# Multi-Dimensional Immuno-Profiling of *Drosophila* Hemocytes by Single Cell Mass Cytometry

**DOI:** 10.1101/2020.06.10.144584

**Authors:** József Á. Balog, Viktor Honti, Éva Kurucz, Beáta Kari, László G. Puskás, István Andó, Gábor J. Szebeni

## Abstract

Single cell mass cytometry (SCMC) combines features of traditional flow cytometry (FACS) with mass spectrometry and allows the measurement of several parameters at the single cell level, thus permitting a complex analysis of biological regulatory mechanisms. We optimized this platform to analyze the cellular elements, the hemocytes, of the *Drosophila* innate immune system. We have metal-conjugated six antibodies against cell surface antigens (H2, H3, H18, L1, L4, P1), against two intracellular antigens (3A5, L2) and one anti-IgM for the detection of L6 surface antigen, as well as one anti-GFP for the detection of crystal cells in the immune induced samples. We investigated the antigen expression profile of single cells and hemocyte populations in naive, in immune induced states, in tumorous mutants (*hop^Tum^* bearing a driver mutation and *l(3)mbn^1^* carrying deficiency of a tumor suppressor) as well as in stem cell maintenance defective *hdc*^*Δ84*^ mutant larvae. Multidimensional analysis enabled the discrimination of the functionally different major hemocyte subsets, lamellocytes, plasmatocytes, crystal cell, and delineated the unique immunophenotype of the mutants. We have identified sub-populations of L2+/P1+ (*l(3)mbn^1^)*, L2+/L4+/P1+ (*hop^Tum^)* transitional phenotype cells in the tumorous strains and a sub-population of L4+/P1+ cells upon immune induction. Our results demonstrated for the first time, that mass cytometry, a recent single cell technology combined with multidimensional bioinformatic analysis represents a versatile and powerful tool to deeply analyze at protein level the regulation of cell mediated immunity of *Drosophila*.

## Introduction

In the animal kingdom, insects have multi-layered innate immune defence mechanisms against invading pathogens. Work on insects, including the fruit fly, *Drosophila melanogaster* which lacks an acquired immune response, plays an important role in our understanding of how innate immunity works [1, 2]. The conserved signaling pathways between insects and vertebrates, combined with the powerful genetic resources provided by *Drosophila*, make this organism an ideal system to model biological phenomena related to human biology and medicine. In *Dorosphila*, microbial infection induces a powerful humoral immune response, the release of antimicrobial peptides, the regulation of which is now well understood [3]. Infection by parasites, development of tumours or wounding induce a cellular immune response by blood cells, the hemocytes, which are capable of sophisticated functions, as recognition, encapsulation and killing of parasites and phagocytosis of microorganisms [4–6]. These functions are exerted by specialized blood cells the phagocytic plasmatocytes, the encapsulating lamellocytes and the melanizing crystal cells. For the identification and characterization of the mechanisms of cell mediated immunity through which the immune cells and tissues can be specifically studied and manipulated, quantitative methods are required. For the definition of the functional hemocyte subsets transgenic reporter constructs and monoclonal antibodies have been developed. These systems generally use fluorescent molecules in the form of *in vivo* markers and antibodies, the use of which significantly contributed to our understanding of innate immunity [7–9]. Recently, single cell mass cytometry was developed to monitor the expression of marker molecules in haematological and other pathological conditions [10,11]. The antibodies against cell type specific antigens are applicable to monitor blood cell differentiation during ontogenesis or following immune induction. However, traditional antibody staining against only one or two of the cell type specific antigens is not sufficient to describe individual hemocyte populations with complex antigen expression patterns. Therefore, we adopted and optimized single cell mass cytometry for *Drosophila* by multiplex analysis of antibodies to transmembrane proteins and intracellular antigens of IgG and IgM type, routinely used for detecting and discriminating hemocyte subsets of *Drosophila melanogaster* [7, 12–16].

The circulating hemocytes of the *Drosophila* larva are classified into three categories, of which only two cell types are present in naive condition. These are the small round phagocytic plasmatocytes, which account for 95% of the circulating hemocytes, and the melanizing crystal cells, which are similar in size to plasmatocytes, but contain prophenoloxidase crystals in their cytoplasm. The third cell type, the large flattened lamellocytes differentiate only in tumorous larvae and in case of immune induction, such as wounding or parasitic wasp infestation [17]. Lamellocytes, together with plasmatocytes are capable of forming a multilayer capsule around the wasp egg, thereby killing the invader [18–20]. Plasmatocytes, crystal cells and lamellocytes can be distinguished with cell type specific monoclonal antibodies, and *in vivo* transgenic reporters [7–9, 12–15]. All plasmatocytes express the P1 antigen (coded by the *nimC1* gene) [21], while lamellocytes show a characteristic expression of L1 (the product of the *atilla* gene), L2, L4, and L6 [14]. Following immune induction, a portion of plasmatocytes transdifferentiate into lamellocytes to fight the parasitic wasp egg [22–25]. This transdifferentiation is accompanied by a stepwise alteration of lamellocyte specific antigen expression.

Understanding cancer, a devastating disease of multicellular organisms is a challenge for scientists. The conserved signal transduction pathways in *Drosophila* with mammals and the easy genetic manipulation made the fruit fly a frequently used model organism to study cancer [26]. Therefore, we investigated two different tumorous *Drosophila* strains, one bearing a driver mutation (*hop^Tum^)* and one carrying deficiency of a tumor suppressor (*l(3)mbn^1^)*. Constitutive activation of the *Drosophila* Janus kinase namely, the Hopscotch (Hop) causes melanotic tumors, lymph gland hypertrophy in the larvae and malignant neoplasia of *hop^Tum^ Drosophila* blood cells [27]. The homozygously mutated state of the tumor suppressor gene, called *lethal* (3) *malignant blood neoplasm* causes malignant transformation, enhanced hemocyte proliferation and lamellocyte differentiation of *l(3)mbn^1^ Drosophila* blood cells [28]. We also investigated the immunophenotype of the mutation of the *hdc* gene (*hdc*^*Δ84*^), which encodes for the *Drosophila* homolog (Headcase) of the human tumor suppressor HECA and plays a role in hematopoietic stem cell maintenance [29, 30]. Wild type *Oregon-R* (*Ore-R*) and *white* mutant *w^1118^* were used as reference strains because *w^1118^* was considered previously as wild type and used for the generation of mutants [31]. Immune activation was monitored successfully by infestation with the *Leptopilina boulardi* parasitoid wasp of *Drosophila* larvae in the *lozenge*>GFP strain *(lz-Gal4, UAS-GFP;* +; +), in which crystal cells were monitored by metal tag labeled anti-GFP antibody [32, 33].

We are the first to demonstrate that single cell mass cytometry is a powerful tool for the characterization of hemocytes in different mutants of *Drosophila* strains at protein level. Bioinformatic analysis revealed the characteristic protein expression pattern of hemocyte subsets at single cell resolution from the studied different genetic variants. These together suggest that single cell mass cytometry is a valuable tool for characterizing immune phenotypes in any model organism, in which antibodies against immune components are available.

## Results and Discussion

### Single cell mass cytometry revealed the transitional phenotypes of hemocytes in the tumorous hop^*Tum*^ and *l(3)mbn^1^*strains

We have built the metal tag labelled panel of discriminative antibodies recognizing *Drosophila melanogaster* hemocytes and hemocyte subsets for mass cytometry. We have conjugated six antibodies against cell surface antigens (H2, H3, H18, L1, L4, P1), against two intracellular antigens (3A5, L2) and one anti-IgM for the detection of L6 surface antigen. List of the antibodies can be found in Table 1. The H18 and 3A5 antibodies reported herein first were characterized and validated before the study by indirect immunofluorescence and Western-blot analysis (Figure S1 and S2). The analysis revealed that 3A5 molecule is expressed in plasmatocytes and lamellocytes in *l(3)mbn^1^*, but not expressed in lamellocytes of immune (*L.b*.) induced larvae (Figure S1), while H18 molecule as a pan-hemocyte marker is expressed in all tested samples (Figure S2). To test and optimize the reactions of the antibodies, a comparative analysis was carried out by correlating the fluorescence activated cell sorting (FACS) (Figure S3A) and the mass cytometry histograms (Figure S3B). The comparison showed similar reactivity patterns. Hemese (H2) pan-hemocyte marker positive single live cells were gated for mass cytometry analysis (Figure S4). All metal-tag labelled antibodies were titrated for mass cytometry as shown in Figure S5. Next, we compared the expansion of the hemocyte populations in the mutants in relation to the two wild type *Ore-R* and *w^1118^*. The proportion of hemocytes expressing the investigated markers were similar in wild type (wt) *Ore-R* and *w^1118^*. However, we detected a significant proliferation of hemocytes expressing the L1, L2, and L4 markers in *l(3)mbn^1^* and *hop^Tum^* mutant larvae, reflecting an extensive differentiation of lamellocytes, a phenotypic characteristic to the blood cell malignancy. A slight elevation in the proportion of L6 expressing hemocytes was also detected (Figure S6 and **Figure 1A**). The explanation for this moderate change may be the fact that L6 is only expressed by a subset of lamellocytes in tumorous larvae [14]. All lamellocyte markers showed a higher expression level in the tumorous *hop^Tum^* mutant compared to the control (Figure S7 and **Figure 1B**). In the *hdc*^Δ*84*^ mutant larvae, we detected a moderate elevation in the expression level of L2, and a decrease in the expression level of P1 (Figure 1B), however, the number of hemocytes expressing lamellocyte markers did not show a significant increase compared to the controls (Figure 1A). This is in line with the finding that in the *hdc*^Δ*84*^ mutant larvae, lamellocytes differentiate in low numbers, while the number of plasmatocytes are reduced [30]. This reduction of plasmatocyte number is also observable in Figure 1A.

**Table 1.** List of the antibodies used for mass cytometry.

| **Marker** | **Clone** | **Isotype** | **Metal tag** | **References** |
| --- | --- | --- | --- | --- |
| H2 (Hemese) | 1.2 | mouse IgG2a | 147 Sm | 12, 14 |
| H3 | 4A12 | mouse IgG1 | 155 Gd | 14 |
| H18 (Tetraspannin42Ed) | H18 | mouse IgG1 | 164 Dy | - |
| L1 (Atilla) | H10 | mouse IgG1 | 149 Sm | 14, 15, 23 |
| L2 | 31A4 | mouse IgG2a | 158 Gd | 14, 23 |
| L4 (Integrin beta-PS) | 1F12 | mouse IgG1 | 159 Tb | 14, 23 |
| L6 (IgM) | H3 | mouse IgM | – | 14, 23 |
| anti-IgM | RMM-1 | rat IgG2a | 172 Yb | 42 |
| P1 (NimC1) | N47 | mouse IgG1 | 154 Sm | 13, 14, 21 |
| 3A5 | 3A5 | mouse IgG2b | 169 Tm | - |
| anti-GFP | – | rabbit<br>polyclonal IgG | 175 Lu | 43 |
| anti-CD45 | HI30 | mouse IgG1 | 89 Y | 44 |

**Figure 1.**
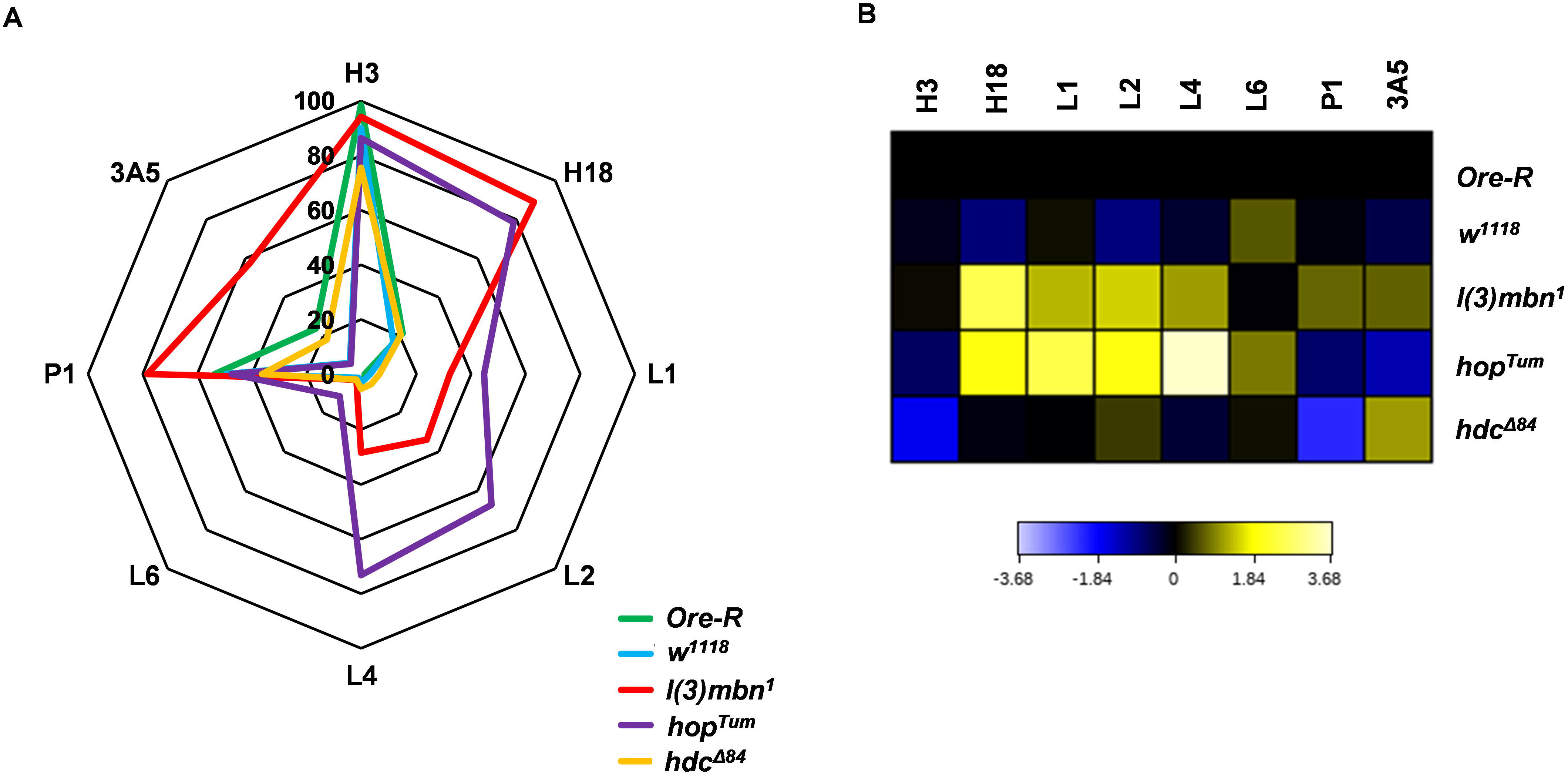
Single cell mass cytometry revealed the expansion of hemocytes in *hop^Tum^* and *l(3)mbn^1^*. **(A)** The percentage of H3, H18, L1, L2, L4, L6, P1, and 3A5 cells were plotted on radar plots for *Drosophila* mutants on *Ore-R* or *w^1118^* background. (**B**) Comparative heatmap of mass cytometry data (arcsinh-transformed median intensity values) regarding marker density at single cell resolution show increased expression of H18, L1, L2, L4 markers in the mutant *hop^Tum^* and *l(3)mbn^1^* in relation to control, the wild type *Ore-R*. Analysis was performed within the H2 (Hemese) positive live singlets.

Multidimensional analysis by the algorithm of t-distributed stochastic neighbor embedding (tSNE) and the visualization of stochastic neighbor embedding (viSNE) was carried out within the H2 (Hemese) positive live singlets based on H3, H18, L1, L2, L4, L6, P1, and 3A5 marker expression in order to map high parametric single cell data on biaxial plots [34]. The viSNE patterns of hemocyte marker expression correlated to the data shown in Figure 1 (**Figure 2**). The viSNE bioinformatic analysis revealed the characteristic protein expression pattern of hemocyte subsets at single cell resolution from the studied genetic variants. We observed a dramatic difference in the viSNE patterns between hemocytes isolated from the tumorous *l(3)mbn^1^* and *hop^Tum^* larvae as compared to either control *Ore-R* or *w^1118^* larvae (Figure 2). Control *Ore-R* or *w^1118^* hemocytes were not discriminated on the viSNE plots showing their minimal genetic distance but tumorous *l(3)mbn^1^* and *hop^Tum^* larvae delineated viSNE maps with the expansion of lamellocytes (Figure 2). In the *hdc*^Δ*84*^ larvae, we detected a subset of hemocytes that express the 3A5 marker at a high level. This subset was detected neither in the control, nor in the tumorous larvae, and may represent a cell type that differentiate as a precursor for lamellocytes as a consequence of the defect in the maintenance of the hematopoietic niche [30].

**Figure 2.**
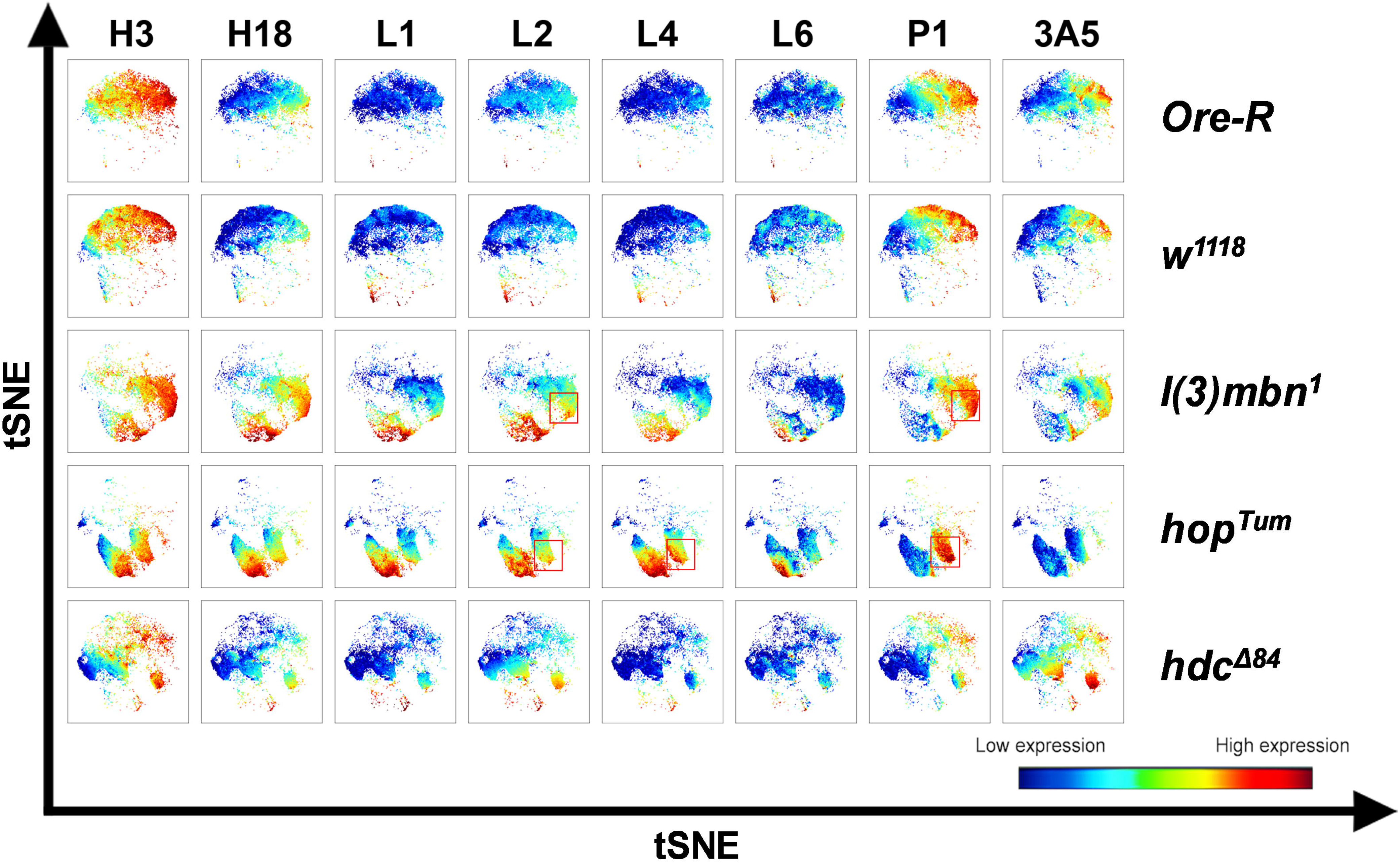
Multidimensional comparative analysis by the tSNE algorithm dissects the cell relatedness of 5 different *Drosophila* strains, namely *Ore-R, w^1118^, l(3)mbn^1^, hop^Tum^* and *hdc*^*Δ84*^. The wild type *Ore-R* and *white* mutant *w^1118^* (genetic backgrounds) are overlapping while both tumorous strains *l(3)mbn^1^* and *hop^Tum^* represent H18, L1, L2, L4 expansion. The tSNE analysis of H3, H18, L1, L2, L4, L6, P1, and 3A5 markers was carried out within the population of pan-hemocyte H2 (Hemese) positive live singlets and visualised as viSNE plots. Subpopulations of cells with common marker expression patterns are grouped close in the multidimensional space, while cells with different marker expression are plotted separately. Coloration is proportional with the intensity of the expression of a given marker: the hotter the plot, the higher the level of expression (red plots). Red boxes mark transitional phenotypes expressing both lamellocyte (L2 or L4) and plasmatocyte (P1) markers.

The Uniform Manifold Approximation and Projection (UMAP) analysis was performed by the hemocyte subset specific, discriminating markers: L1, L2, L4, L6 for lamellocytes and P1 for plasmatocytes on the 5 studied genetic variants of *Drosophila melanogaster*. The UMAP analysis resulted in the same conclusion as tSNE, namely, that lamellocyte expansion occurs in in tumorous strains *l(3)mbn^1^* and *hop^Tum^* (Figure S8). Both the viSNE and UMAP analysis demonstrate transitional phenotypes of certain lamellocytes and plasmatocytes by the transitional coloration of marker expression (partially overlapping L2+ or L4+ with some P1+ cells) at protein level in *l(3)mbn^1^* and *hop^Tum^*. Merging viSNE graphs outlined characteristic maps of each strain based on high parametric mass cytometry data (**Figure 3A-C**). The *Ore-R* and *w^1118^* controls showed overlapping patterns on the viSNE diagram (Figure 3A-C), with a somewhat lower expression of all markers observed in case of the *w^1118^*, which may be due to uncontrollable genetic background variations. The dots representing to *hdc*^Δ*84*^ hemocytes, mutant of the *hdc* regulator of hematopoietic stem cell maintenance [30], were detected as a zone in between the control and the tumorous patterns (Figure 3C). The most likely explanation to this phenomenon is that *hdc*^Δ*84*^ homozygous larvae produce lamellocytes, but in a much lower proportion than tumorous larvae, the *l(3)mbn^1^* and *hop^Tum^* [30]. Tumorous hemocytes *l(3)mbn^1^* and *hop^Tum^* were closely mapped and partially overlapping, giving a population clearly separated from the cloud of the controls, due to the lamellocye-expansive malignant phenotype (Figure 3A-C).

**Figure 3.**
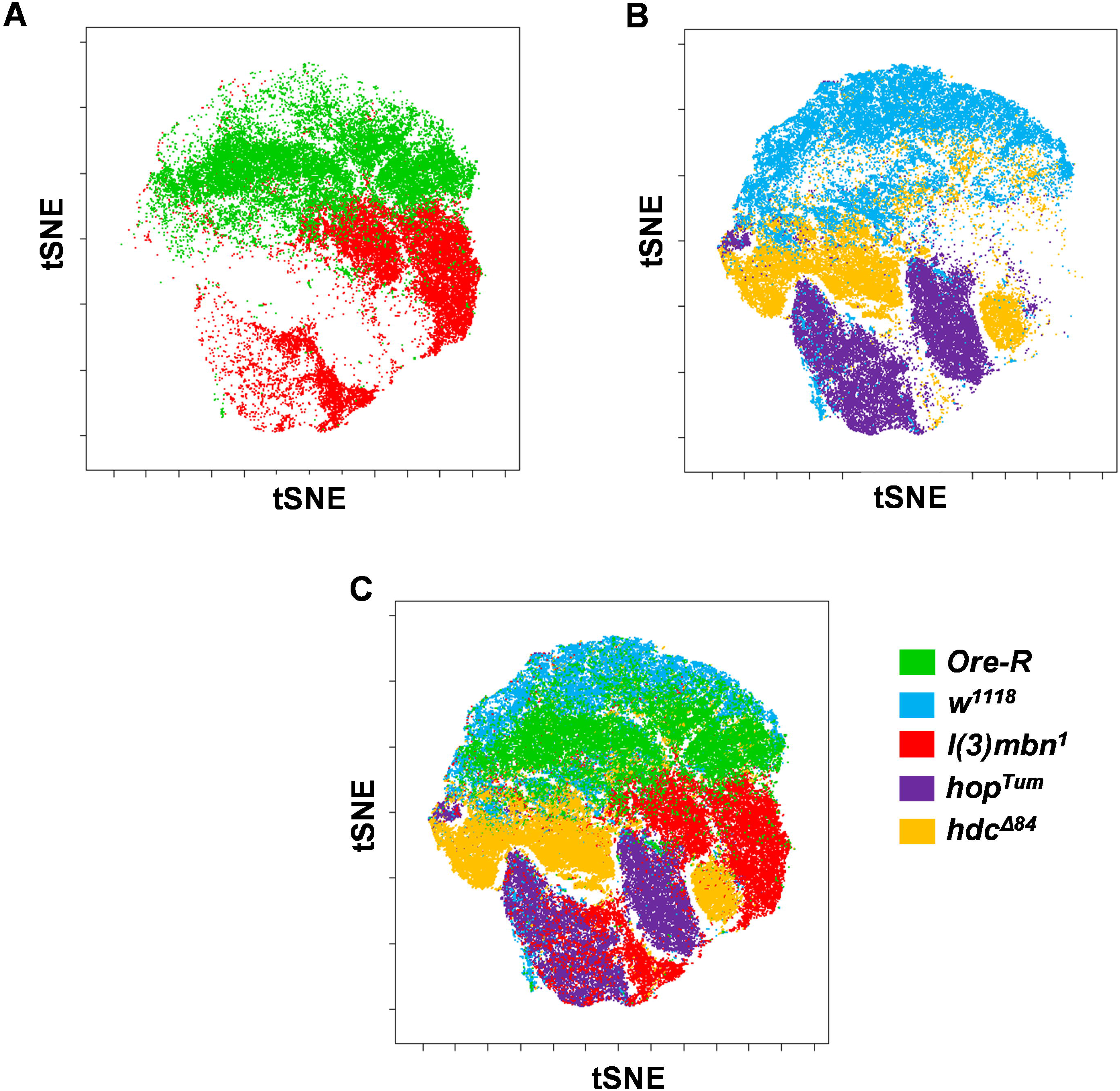
Merging viSNE graphs (based on H3, H18, L1, L2, L4, L6, P1, and 3A5 marker expression within the pan-hemocyte H2 (Hemese) positive live singlets) outlines characteristic maps of each strain (green = *Ore-R*, blue = *w^1118^*, red = *l(3)mbn^1^*, lilac = *hop^Tum^*, yellow = *hdc^Δ84^*) based on high parametric mass cytometry data. (**A**) The viSNE comparison of *l(3)mbn^1^* and its wt counterpart, the *Ore-R*. (**B**) The viSNE comparison of *w^1118^*, *hop^Tum^,* and *hdc*^*Δ84*^. (**C**) The viSNE islands of the control cells (*Ore-R* and *w^1118^)* localize separately from the tumorous *l(3)mbn^1^* and *hop^Tum^* hemocytes while *hdc*^*Δ84*^ represents a transition phenotype.

### Single cell mass cytometry revealed the transitional phenotypes of hemocytes upon immune induction

In order to monitor the changes in the composition of hemocyte subsets following immune induction, we used *lz>GFP* larvae and complemented the experiment with anti-GFP labeling, which enables the detection of crystal cells [32, 33]. The tSNE analysis of H3, H18, L1, L2, L4, L6, P1, 3A5 markers and anti-GFP (marking crystal cells in this particular system) was carried out within the population of pan-hemocyte H2 (Hemese) positive live singlets (**Figure 4A**). We observed a new subset of hemocytes appearing 72 h after infestation of the *lz>GFP* larvae with the parasitic wasp (Figure S9. and Figure 4A). This subset of cells accounts for the lamellocytes that differentiate as a result of the immune induction, since these cells fall into the high expression part of the viSNE for the L1, L2, L4, and L6 lamellocyte markers (Figure S9. and Figure 4A). This finding is in correlation with the increase of the number of hemocytes expressing the L1 (35.1% vs. 1.81%), L2 (32% vs. 1.6%), L4 (34.36% vs. 1.39%) and L6 (13.82 vs. 0.935%) markers (**Figure 4B**), and the elevated expression levels of the lamellocyte markers detected in immune induced larvae compared to the naive control (**Figure 4C**). Interestingly, a new subset of crystal cells (anti-GFP ^+^ cells) also appeared in immune induced (*lz>GFP i.i.)* larvae compared to the control (*lz>GFP)* (Figure 4A). The viSNE pattern of the 3A5 marker also changed significantly after the immune induction, which may be due to the newly differentiating hemocytes, similarly to that observed in the *hdc*^Δ*84*^ larvae (Figure 4A). Taken together, we report herein the first panel of metal-conjugated anti-*Drosophila* antibodies to present the applicability of mass cytometry for that canonical model organism of genetics. Recent studies identified novel subpopulations of *Drosophila* hemocytes based on single cell RNA data [35–38]. These findings largely contributed to the definition of hemocyte clusters and to the characterization of intermediate cells in the transition from plasmatocyte to lamellocyte. In these experiments, clusters were defined by the gene expression patterns of individual hemocytes. The application of CyTOF (cytometry by time-of-flight) can complement these comprehensive transcriptomic studies and verify the existence of transitional phenotypes at protein level. The comparative analysis of *Ore-R* and *white^1118^* with *l(3)mbn^1^*, *Hop^Tum^*, *hdc*^*Δ83*^ revealed transitional phenotypes at protein level and the differences among reference stains: *Ore-R* and *white^1118^*. Both the viSNE and UMAP analysis demonstrated transitional phenotype of certain sub-populations of lamellocytes and plasmatocytes by the transitional coloration of common marker expression (partially overlapping L2+ or L4+ with P1+ cells) at protein level in *l(3)mbn^1^*, *hop^Tum^*. This has been verified by a functional assay of immune induction (Figure 4). Our study demonstrates transitional phenotypes (Figure 2, Figure 4, Figure S8) from single cell data at protein level which places the innate immunity of *Drosophila* in a new biological insight. Additionally, we report herein two novel hemocyte markers, H18 located on the cell surface and 3A5 with intracellular localization. The simultaneous detection of several antigens provided by CyTOF could not be achieved earlier by traditional microscopy. The main advantage of CyTOF is the multidimensionality coupled with complex computational tools, therefore we propose the extension of the basic panel used in our study with antibodies recognizing signaling molecules (e.g. MAP kinases), enzymes (to follow metabolic pathways), cellular structural proteins (e.g. cytoskeletal, cargo proteins) up to 42 markers in one single tube. Another advantage of the presented method is that CyTOF enables investigations at protein level (data of transcriptomics should be also verified at protein level) with single cell resolution. However, we may consider the limitations of the CyTOF which are a.) the availability of antibodies against the protein of interest (which is also a limitation for other antibody-based detection approaches). Moreover, anti-tag antibodies are available when the protein of interest is labelled with a fusion tag, or the cell of interest is labelled with the expression of a marker protein (we report herein the use of anti-GFP). Another limitations are b) the availability of the CyTOF technology (it is increasing and most of the research centres are supposed to own the technology, as there were 94 instruments already installed in Europe in 2020 January), c) the relative high cost of the CyTOF technology (although the cost should be taken into account by the number of investigated markers at protein level and the number of single cells).

**Figure 4.**
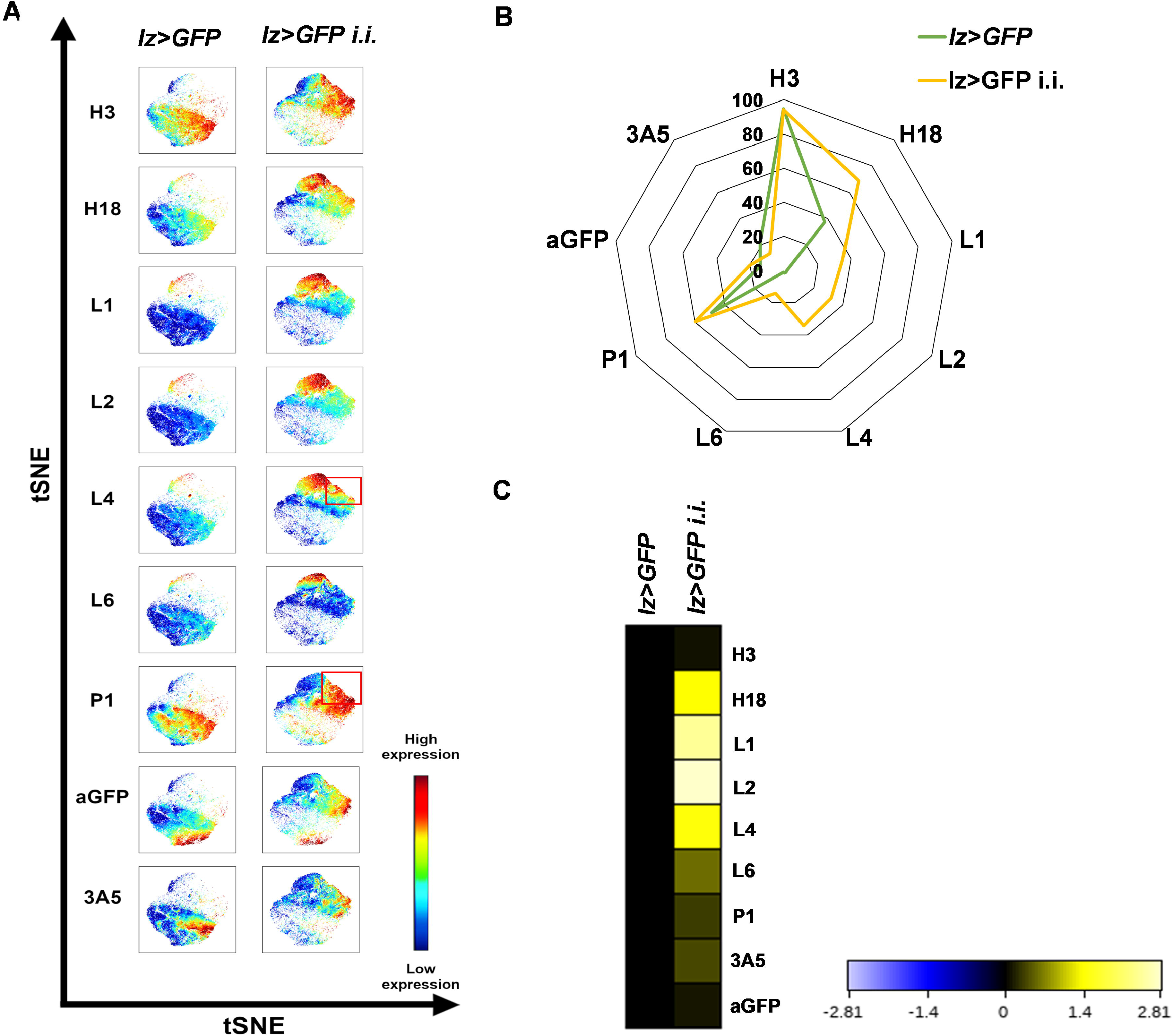
Immune activation was monitored successfully by infestation with the *Leptopilina boulardi* parasitoid wasp of the *lozenge*>*GFP* strain. (**A**) viSNE analysis of naive (*lz*>*GFP*) and immune induced (*lz*>*GFP i.i.*) *Drosophila* larvae. The tSNE analysis of H3, H18, L1, L2, L4, L6, P1, 3A5 markers and anti-GFP (marking crystal cells in this particular system) was carried out within the population of pan-hemocyte H2 (Hemese) positive live singlets. Red boxes mark a subpopulation, the transitional phenotype of hemocytes expressing both lamellocyte (L4+) and plasmatocyte (P1) markers upon immune induction. (**B**) The percentage of H3, H18, L1, L2, L4, L6, P1, anti-GFP (crystal cells), and 3A5 positive cells. (**C**) The heatmap of the (arcsinh-transformed) median values shows the expression changes of the hemocyte marker expression upon immune induction. Analysis was performed within the pan-hemocyte marker H2 (Hemese) positive live singlets.

We believe that our method serves as a rapid and cost-effective tool to monitor the alteration of hemocyte composition influenced by various agents or mutations. In those cases, it is less expensive and easier to perform than single-cell transcriptome analysis. Additionally, the CyTOF can complement transcriptomic studies verifying up to 42 simultaneous markers at protein level with single cell resolution.

## Conclusion

The SCMC combines the features of traditional cytometry with mass spectrometry and enables the detection of several parameters at single cell resolution, thus permitting a complex analysis of biological regulatory mechanisms. We optimized this platform to analyze the cellular elements, the hemocytes of the *Drosophila* innate immune system. The SCMC analysis with 9 antibodies to all hemocytes and hemocyte subsets showed a good accordance of fluorescence flow cytometry results, in terms of positivity on hemocytes of the tumor suppressor mutant *l(3)mbn^1^*. Further, we investigated the antigen expression profile of single cells and hemocyte populations in *Ore-R and w^1118^* controls, and tumorous (*l(3)mbn^1^*, *hop^Tum^*) strains, as well as in a stem cell maintenance defective mutant (*hdc*^*Δ84*^). The immunophenotype of immune activation upon infestation with a parasitoid wasp, the differentiation of lamellocytes was detected by 10 antibodies in the *lz>GFP*.

Multidimensional analysis (viSNE) enabled the discrimination of the major hemocytes: lamellocytes, plasmatocytes, crystal cells and delineated the unique single cell immunophenotype of the mutant strains under investigation. Single cell mass cytometry identified sub-populations of L2+/P1+ (*l(3)mbn^1^)*, L2+/L4+/P1+ (*hop^Tum^)* transitional phenotype cells in the tumorous strains and a sub-population of L4+/P1+ cells upon immune induction. We demonstrated that mass cytometry, a recent single cell technology coupled with multidimensional bioinformatic analysis at protein level represents a powerful tool to deeply analyze *Drosophila*, a key multicellular model organism of genetic studies with a wide inventory of available mutants.

## Materials and methods

### *Drosophila* stocks

The following *Drosophila* lines were used in the study: *w^1118^* (BSC#9505), *ORE-R* (wild type), *w; hdc*^*Δ84*^*/TM3, Kr>GFP* [30], *lz-Gal4, UAS-GFP;* +; + (a gift from Bruno Lemaitre, Lausanne, Switzerland) [32], *l(3)mbn^1^/TM6 Tb* [28], a homozygous *hop^Tum^* (BSC#8492) line generated by dr. Gábor Csordás (BRC, Szeged, Hungary). The flies were grown on a standard cornmeal-yeast substrate at 25 °C.

### Production of the H18 and 3A5 antibodies

Monoclonal antibodies against *Drosophila* hemocytes were raised as described previously [14]. Briefly, BALB/c mice were immunized by i.p. injection of 10^6^ hemocytes from late third instar larvae of the *lethal(3)malignant blood neoplasm* [*l(3)mbn^1^*] mutant larvae in *Drosophila* Ringer’s solution (Sigma-Aldrich, St. Louis, MI, USA). Booster injections were given 4, 8, and 13 weeks later. Three days after the last immunization, spleen cells were collected and fused with SP2/O myeloma cells by using polyethylene glycol (PEG1450, P5402 Sigma-Aldrich). Hybridomas were selected in HAT medium (HAT = hypoxanthine-aminopterin-thymidine Supplement, 21060017 Thermo Fischer Scientific Waltham, MA, USA) and maintained as described by Kohler and Milstein [14, 39]. Hybridoma culture supernatants were screened by indirect immunofluorescence on acetone fixed, permeabilized and on live hemocytes. The selected hybridomas were subcloned three times by limiting dilution.

### Isolation of hemocytes

Hemocytes were isolated from late third stage larvae by dissecting the larvae in *Drosophila* Schneider’s solution (21720001 Thermo Fisher Scientific, Waltham, MA, USA)) supplemented with 5% fetal bovine serum albumin (FBS, F7524-500ML Sigma-Aldrich) plus 0.003% 1-phenyl-2-thiourea (P7629 Sigma-Aldrich).

### Immune induction

*lz-Gal4; UAS-GFP* flies (*lz>GFP*) laid eggs for three days in bottles containing standard *Drosophila* medium. After 72 hours, larvae were infected with *Leptopilina boulardi* wasps for 6 hours. Larvae with visible melanotic nodules were selected 72 hours after infestation for isolation of hemocytes. Age and size-matched larvae were used as control.

### Immunofluorescent staining

Immunofluorescent staining was performed as described previously [23]. Briefly, hemocytes were attached to multispot slides (SM-011, Hendley-Essex, Loughton, UK) at 21 °C for 45 min. Fixation was performed with acetone for 6 min, rehydrated and subsequently blocked for 20 min in PBS supplemented with 0.1% BSA (PBS = phosphate buffered saline, P4417 Sigma-Aldrich; BSA = bovine serum albumin, A2058 Sigma-Aldrich), incubated with the indicated antibodies for 1 h at 21 °C, washed three times with PBS and incubated with CF-568 conjugated anti-mouse IgG (H+L), F(ab’)2 fragment (1:1000, SAB4600082 Sigma-Aldrich) for 45 min. Nuclei were labeled with DAPI (D9542 Sigma-Aldrich). The microscopic analysis was carried out using a Zeiss Axioskope 2MOT epifluorescent microscope and Axiovision 2.4 software (Zeiss, Oberkochen, Germany).

### Western blotting

Western blotting was performed in order to test the specificity of the anti-3A5 and anti-H18 antibodies as described previously [12]. Briefly, proteins were differentiated by SDS-PAGE. Following the electrophoresis, the proteins were blotted onto nitrocellulose membrane (Hybond-C, 10564755 Amersham Pharmacia, Buckinghamshire, UK) in the transfer buffer (25 mM Tris pH 8.3, 192 mM glycine, 20% (V/V) methanol). The nonspecific binding was blocked with PBS supplemented with 0.1% Tween 20 (PBST, P1379 Sigma-Aldrich) and 5% non-fat dry milk at 21 °C for 1 h. The blotted proteins were reacted to the indicated antibody (anti-3A5 in Figure S1, and anti-H18 in Figure S2) with rotation at 21 °C for 3 h. Washing was performed with PBST three times for 10 min and then incubated with HRPO-conjugated anti-mouse antibody (62-6520 Thermo Fisher Scientific). After three washes with PBST for 10 min, the proteins were detected by the ECL-Plus system (32132 Thermo Fisher Scientific) following the manufacturer’s recommendations.

### Flow cytometry

Flow cytometry was executed as published previously [12]. Briefly, 20 μl of 10^7^/ml hemocyte suspension was plated in insect Schneider’s medium (supplemented with 10% FCS) into each well of a 96-well U-bottom microtiter plate (3635 Corning Life Sciences, Tewksbury, MA, USA). Samples for intracellular staining were treated by 2% paraformaldehyde (158127 Sigma-Aldrich). Hybridoma supernatants (50 μl) were measured to each well, and reacted at 4 °C for 45 min. The negative control monoclonal antibody was a mouse IgG1 (clone T2/48, anti-human anti-CD45) [40]. After the incubation, cells were washed three times with ice-cold Schneider’s medium. The secondary antibody, Alexa Fluor 488-labeled anti-mouse IgG (AP124JA4 Sigma-Aldrich) was added (1:1000). After 45 min incubation at 4 °C, the cells were washed (three times) with ice-cold Schneider’s medium and acquired on FACSCalibur (Beckton Dickinson, Franklin Lakes, NJ, USA).

### Mass cytometry

Mass cytometry was performed as we published earlier with some modifications [10, 41]. The affinity purified monoclonal antibodies were provided by Istvan Ando’s group (BRC, Szeged, Hungary) (Table 1) or purchased: anti-IgM, (406527 Biolegend, San Diego, CA, USA [42]), anti-GFP (A11122 Thermo Fisher Scientific [43]), anti-CD45 (3089003B Fluidigm, South San Francisco, CA, USA [44]) and conjugated in house according to the instructions of the manufacturer (Maxpar antibody labeling kit, Fluidigm). Optimal antibody concentrations were titrated prior use (Figure S5). The following antibody concentrations were used: H2: 5 μg/ml, H3: 5 μg/ml, H18: 5 μg/ml, L1: 1 μg/ml, L2: 7.5 μg/ml, L4: 7.5 μg/ml, L6: 10 μg/ml, anti-IgM: 10 μg/ml, P1: 7.5 μg/ml, 3A5: 5 μg/ml, anti-GFP: 10 μg/ml. The negative control monoclonal antibody was a mouse IgG1 (clone Hl30, anti-human 89Y labeled anti-CD45) in 1:100 dilution. The isotypes of anti-*Drosophila* antibodies were determined by the IsoStrip™ Antibody Isotyping Kit (11493027001 Roche, Basel, Switzerland) according to the instructions of the manufacturer.

Single cell suspensions were centrifugated at 1100 g at 6 °C for 4 min and incubated with viability marker (5 μM cisplatin, 195 Pt, 201064 Fluidigm) on ice in 40 μl PBS for 3 min. Cells were washed twice with 200 μl Maxpar Cell Staining Buffer (MCSB, 201068 Fluidigm) and centrifugated at 1100 g at 6°C for 4 min. Cells were resuspended in 50 μl MCSB and 50 μl surface antibody cocktail (2 ×) was added, incubated on ice for 30 min. Cells were washed with 200 μl MCSB and stained with anti-IgM antibody (volumes were the same as in the surface staining), incubated on ice for 30 min. Cells were washed with 200 μl MCSB and suspended in 100 μl 1 × Maxpar Fix I buffer (201065 Fluidigm), incubated on ice for 20 min. Cells were washed twice with 200 μl PermS buffer (201066 Fluidigm) then stained with the intracellular antibody cocktail (L2, 3A5 and anti-GFP in *Lz>GFP* samples), left on ice for 30 min. Cells were washed once with MCSB then fixed with 200 μl 1.6% formaldehyde solution (freshly diluted from 16% Pierce formaldehyde in PBS, 28906 Thermo Fisher Scientific), incubated on ice for 10 minutes then centrifugated at 1300 g at 6°C for 4 min. After fixation, cells were resuspended in 300 μl Maxpar Fix and Perm buffer (201067 Fluidigm) containing 125 nM Cell-ID DNA intercalator (191/193 Iridium, 201192A Fluidigm) and incubated at 4 °C overnight. Before the acquisition samples were washed in MCSB twice and in PBS once (without Mg^2+^ and Ca^2+^, 10010015 Thermo Fisher Scientific) by centrifugation at 1300 g at 6°C for 4 min. Cells were counted using Bürker chamber. For the measurement on Helios, the concentration of cells was set to 0.5 × 10^6^/ml in cell acquisition solution (CAS, 201240 Fluidigm) supplemented with 10% EQ Calibration Beads (201078 Fluidigm). Cells were filtered (30 μm, 04-0042-2316 Celltrics, Sysmex Partec, Görlitz, Germany) prior to acquisition. Samples were run on CyTOF (cytometry by time-of-flight) Helios (Fluidigm). Bead based normalization of CyTOF cytofdata was performed. After randomization, normalization and FCS file generation the files were further analyzed in Cytobank (Beckman Coulter, Brea, CA, USA). Analysis of the cells was carried out on live singlets within the pan-hemocyte marker, H2 positive population. The viSNE (visualization of stochastic neighbour embedding) analysis was carried out on 3 × 10^4^ cisplatin negative (live) singlets with the following settings: iterations = 1000, perplexity = 30, theta = 0.5).

## Authors’ contributions

JAB carried out the mass cytometric experiments, analysis and visualization

VH participated in *Drosophila* work, drafted the manuscript and supervised the analysis

EK produced and affinity purified the antibodies, carried out flow cytometric experiments, prepared graphs and supervised the analysis, and revised the manuscript

LGP supervised the study and revised the manuscript

IA provided the antibodies, supervised the study, and revised the manuscript

GJS designed and supervised the study, designed the experiments and analysis, prepared the figures, drafted the manuscript.

The authors read and approved the final version of the manuscript.

## Acknowledgements

This work was supported by the following grants: GINOP-2.3.2-15-2016-00001, GINOP-2.3.2-15-2016-00030 (LGP), GINOP-2.3.2-15-2016-00035 (ÉK), and NKFI NN118207 and NKFI K120142 (IA), NKFI 120140 (EK), OTKA K-131484 (VH) by the National Research, Development and Innovation Office. Gábor J. Szebeni was supported by the New National Excellence Program of the Ministry for Innovation and Technology (UNKP-19-4-SZTE-36) and by the János Bolyai Research Scholarship of the Hungarian Academy of Sciences (BO/00139/17/8). We are grateful to Mrs. Olga Kovalcsik for the technical help.

