## Supplementary material for "Multi-Dimensional Immuno-Profiling of *Drosophila* Hemocytes by Single Cell Mass Cytometry": SupplFig_legend_bioRxiv.docx

**Figure S1 Expression pattern and molecular weight of the 3A5 marker molecule**

(**A**) The expression of 3A5 in the wild type *Ore-R* and *L. boulardi G486* immune induced *Ore-R* larvae and (**B**) in the *l(3)mbn^1^* tumor suppressor mutant circulating hemocytes. The 3A5 molecule is expressed in plasmatocytes and lamellocytes in *l(3)mbn^1^*, but not expressed in lamellocytes of immune (*L.b*.) induced larvae. Fluorescence microscopy and indirect immunoﬂuorescence analysis were carried out with a Zeiss Axioskope 2 MOT epiﬂuorescence microscope. Images were taken with a Zeiss Axiocam digital camera. Scale bars: 10 µm (**C**) Detection of the 3A5 molecule from *l(3)mbn^1^* hemocyte extract. 3A5 is detected as a 100 kDa protein in Western blot.

**Figure S2 Expression pattern and molecular weight of the H18 marker molecule**

(**A**) The expression of H18 in the wild type *Ore-R* and *L. boulardi G486* immune induced *Ore-R* larvae and (**B**) in the *l(3)mbn^1^* tumor suppressor mutant circulating hemocytes. H18 molecule is expressed in plasmatocytes and lamellocytes both in naive and in immune (*L.b*.) induced larvae in *Ore-R,* and in *l(3)mbn^1^* tumor suppressor mutant circulating hemocytes*.* Fluorescence microscopy and indirect immunoﬂuorescence analysis were carried out with a Zeiss Axioskope 2 MOT epiﬂuorescence microscope. Images were taken with a Zeiss Axiocam digital camera. Scale bars: 10 µm (**C**) Detection of the H18 molecule from *l(3)mbn^1^* hemocyte extract. H18 is detected as a 22.5 kDa protein in Western blot.

**Figure S3 Mass cytometry histograms were verified by conventional fluorescent flow cytometry in Dm hemocytes**

Comparison of (**A**) fluorescence flow cytometry (FACS) data and (**B**) mass cytometry (CyTOF) data obtained from the same *l(3)mbn^1^* hemocytes and same H2, H3, H18, L1, L2, L4, L6, P1, and 3A5 anti-hemocyte antibodies. The negative control monoclonal antibody used for FACS was of mouse origin, the T2/48 IgG1 (anti human anti-CD45). The negative control monoclonal antibody used for CyTOF was of mouse origin, the HI30 IgG1 (anti human anti-CD45) labeled by 89Y metal tag.

**Figure S4 Gating strategy to define cells, singlets, living cells and H2+ pan-hemocytes** Drosophila hemocytes were stained with cisplatin (195 Pt), H2 antibody (147Sm), DNA intercalators (191Ir and 193Ir) as described in the Materials and Methods section. Cells were gated in Cytobank to discriminate cells from calibration beads (DNA 191Ir^+^/140Ce^-^), to define singlets, to determine living cells (cisplatin 195Pt^-^) and H2^+^ hemocytes in *l(3)mbn^1^*strain.

**Figure S5 Titration of the metal-tag labeled antibodies for mass cytometry**

Antibody titration was performed in order to determine the optimal concentration of (**A**) anti-H2, anti-H3, anti-H18, anti-L1, anti-L2, anti-L4, anti-P1, anti-3A5, and (**B**) anti-L6, anti-IgM (concentrations are indicated in the figure) antibodies in *l(3)mbn^1^*strain within the living singlets. The negative control monoclonal antibody used for CyTOF was of mouse origin, the HI30 IgG1 (anti human anti-CD45) labeled by 89Y metal tag. (**C**) The anti-GFP antibody detecting crystal cells was titrated in the *(lz>GFP)* lozenge strain within the H2+ (living singlet) hemocytes.

**Figure S6**  **Representative dot plots of mass cytometry measurements using metal conjugated antibodies**

Antibody labeling was performed in order to detect H3, H18, L1, L2, L4, L6, P1, and 3A5 markers (within H2+ living singlets) in the *Ore-R*, *w^1118^*, *l(3)mbn^1^*, *hop^Tum^*, *hdc^Δ84^* strains as described in Materials and Methods section.

**Figure S7** **Representative histograms of mass cytometry measurements using metal conjugated antibodies**

Antibody labeling was performed in order to detect H3, H18, L1, L2, L4, L6, P1, and 3A5 markers (within H2+ living singlets) in the *Ore-R*, *w^1118^*, *l(3)mbn^1^*, *hop^Tum^*, *hdc^Δ84^* strains as described in Materials and Methods section.

**Figure S8 The Uniform Manifold Approximation and Projection (UMAP) analysis showed that lamellocyte expansion occurs in in tumorous strains *l(3)mbn^1^* and *hop^Tum^***

The UMAP analysis was performed by the hemocyte subset specific, discriminating markers: L1, L2, L4, L6 for lamellocytes and P1 for plasmatocytes on the 5 studied genetic variants of Drosophila melanogaster. The metrics were as follows: Euclidean, nearest neighbors:20; min. distance:0.5; 20000 cells/samples in FlowJo ™ v10.6.1 (Becton Dickinson). Red boxes mark transitional phenotypes of hemocytes expressing both lamellocyte (L2 or L4) or plasmatocyte (P1) markers.

**Figure S9**  **Representative histograms of mass cytometry measurements using metal conjugated antibodies upon activation**

Antibody labeling was performed in order to detect H3, H18, L1, L2, L4, L6, P1, crystal cells (anti-GFP) and 3A5 markers in the naive *(lz>GFP)* and immune induced *(lz>GFP i.i.)* larvae as described in Materials and Methods section.
