## Supplementary figures and images for "Multi-Dimensional Immuno-Profiling of *Drosophila* Hemocytes by Single Cell Mass Cytometry"

### Figure S1.tif

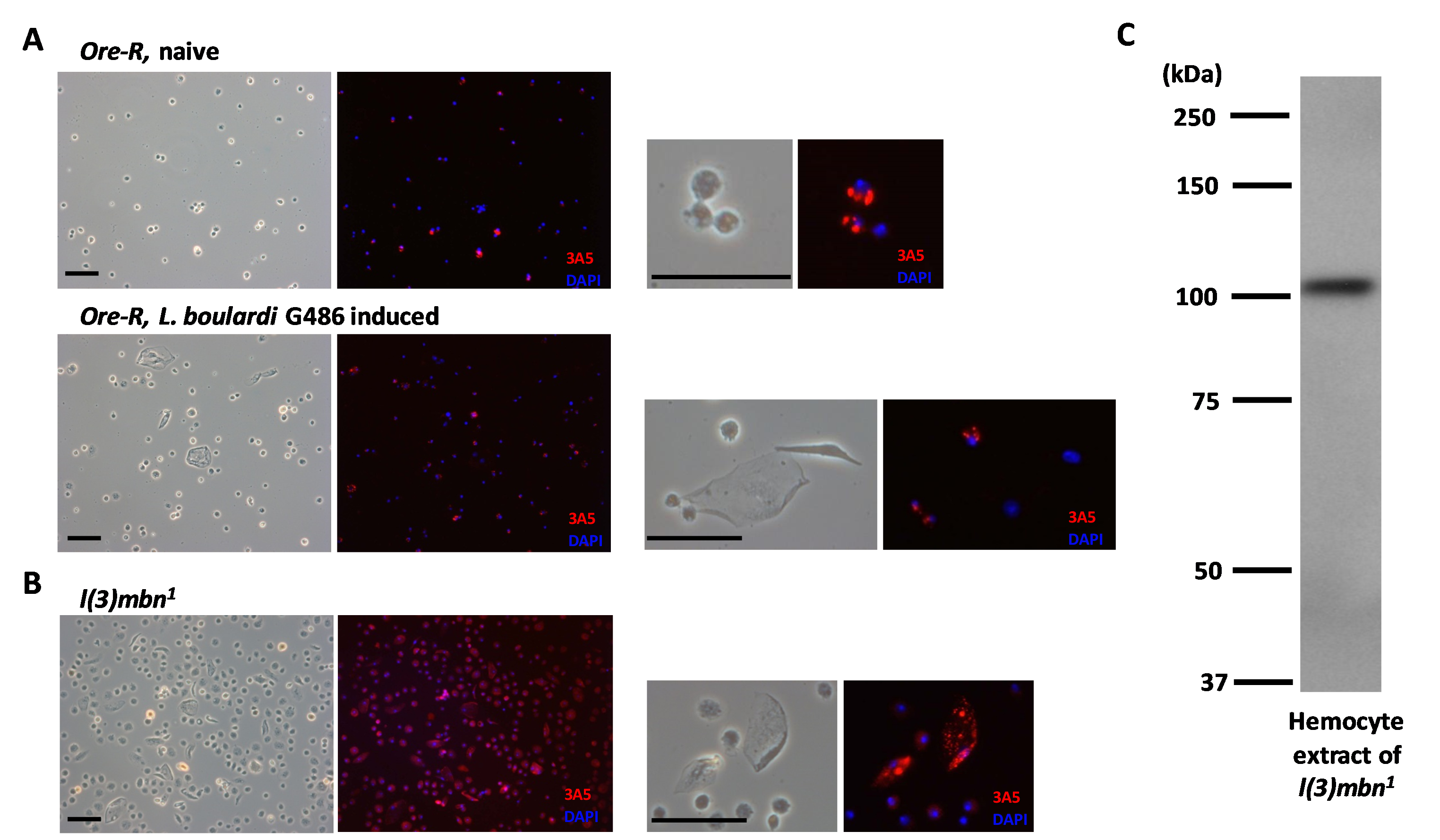

### Figure S2.tif

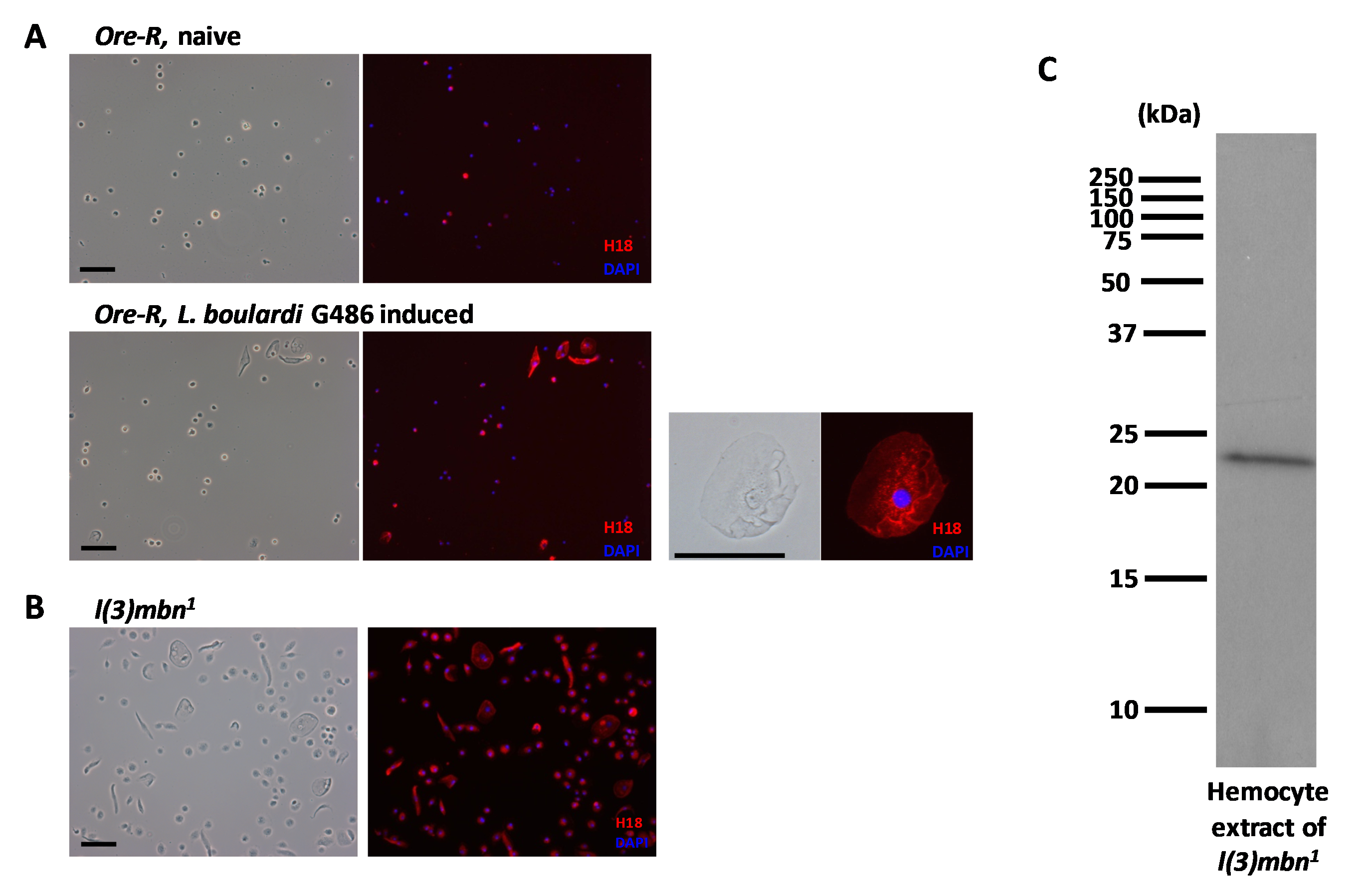

### Figure S3.tif

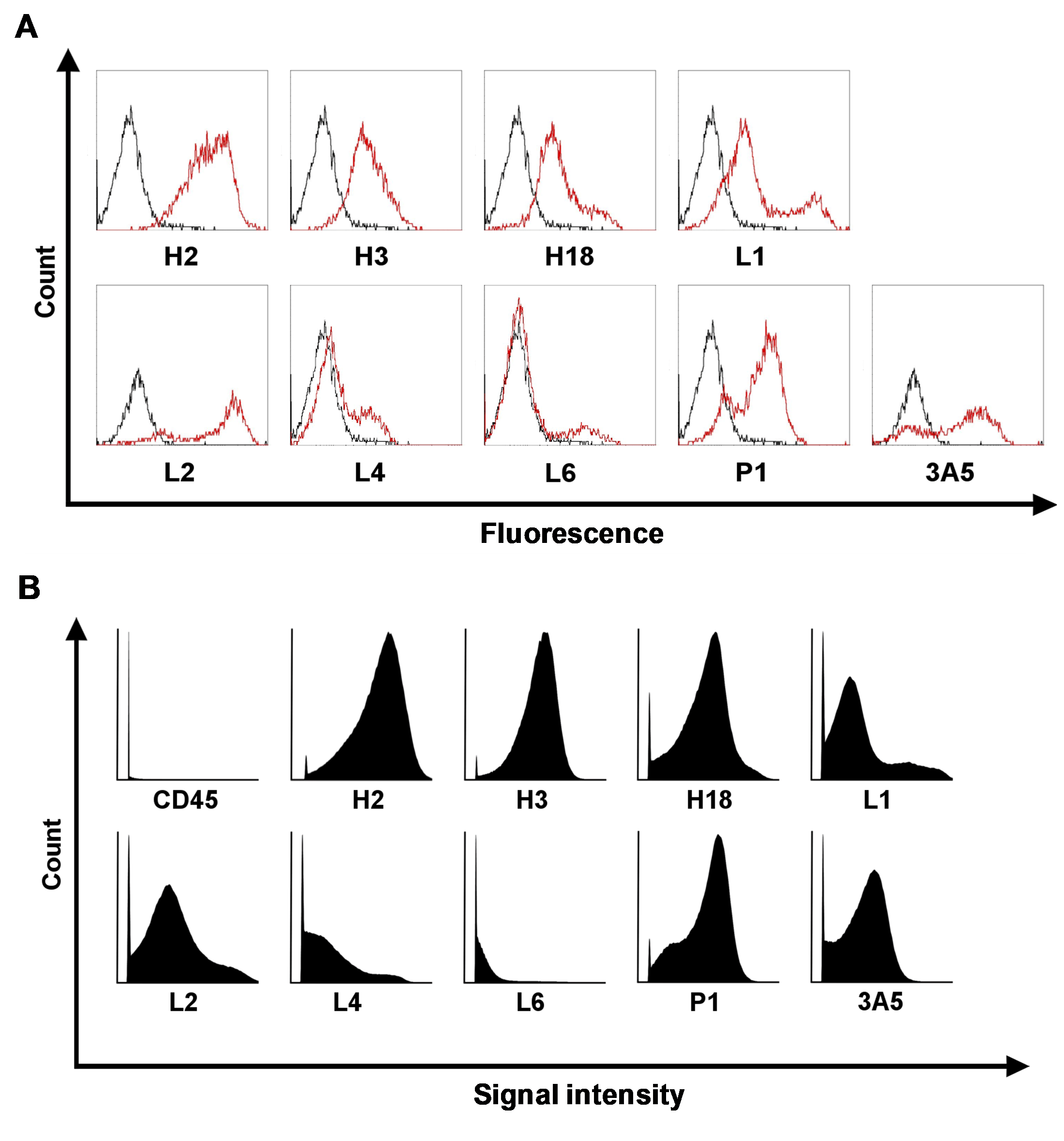

### Figure S4.tif

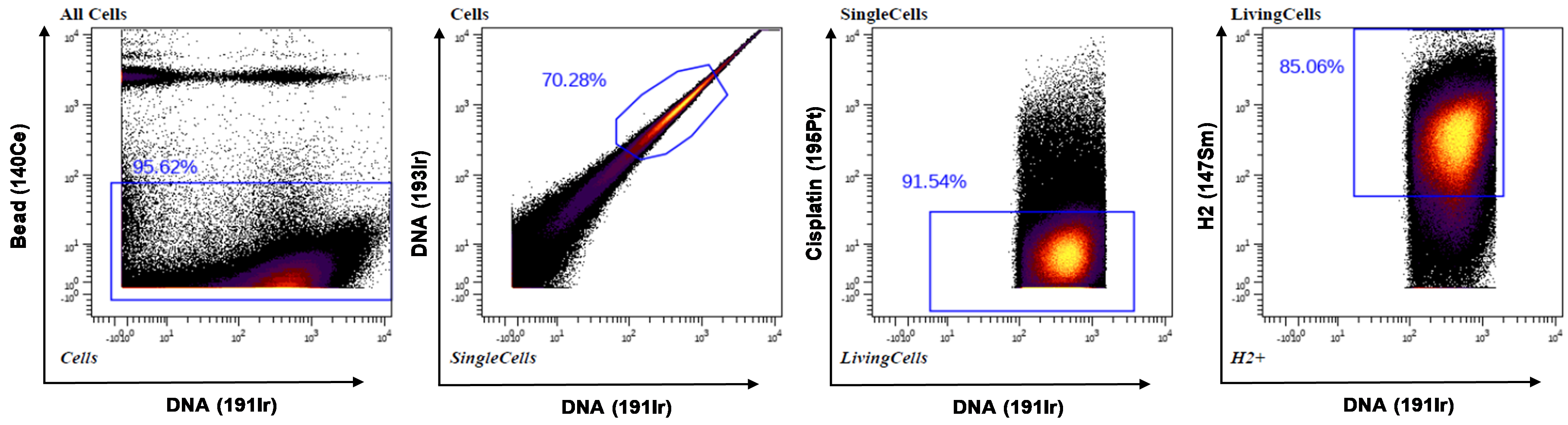

### Figure S5.tif

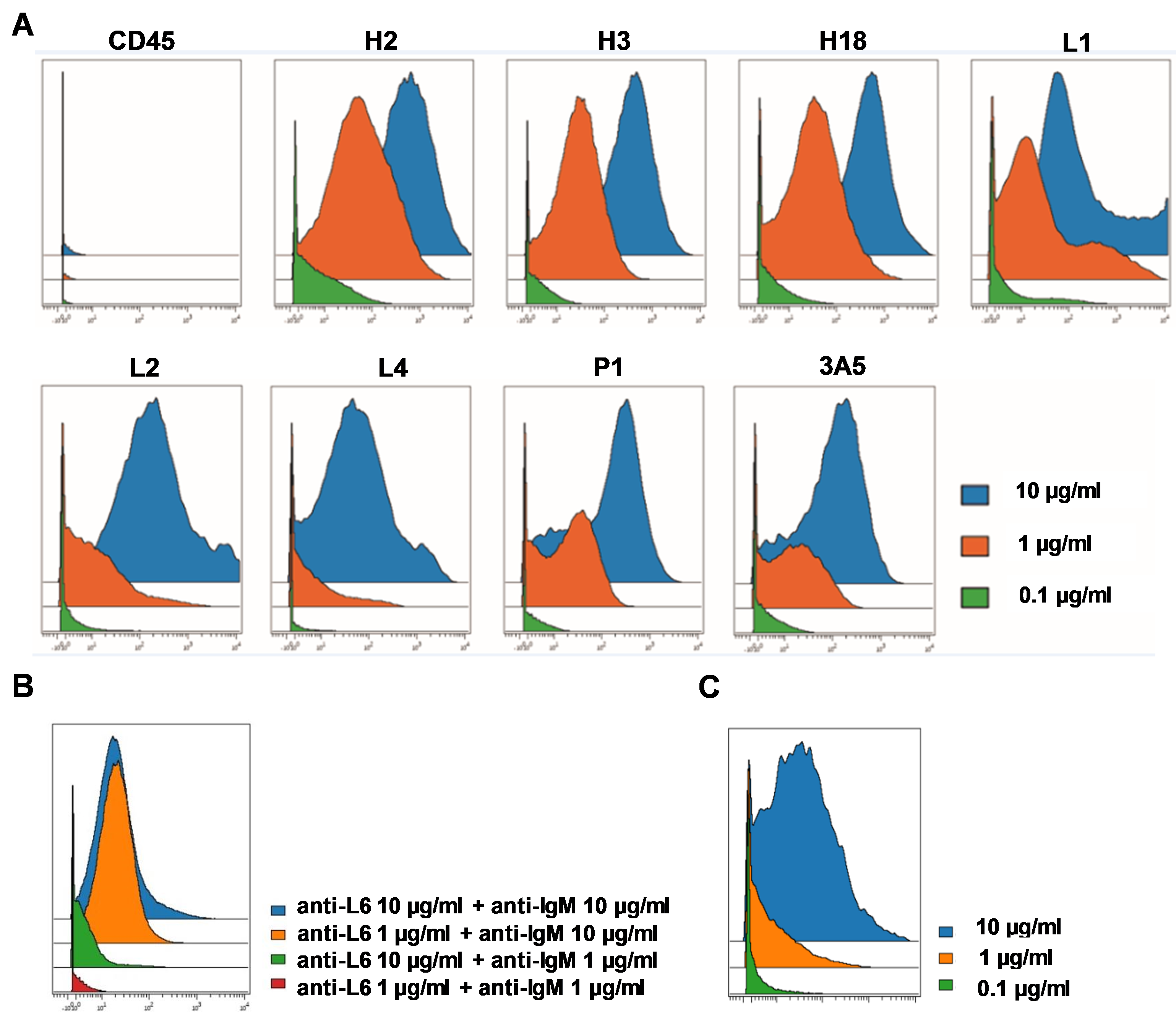

### Figure S6.tif

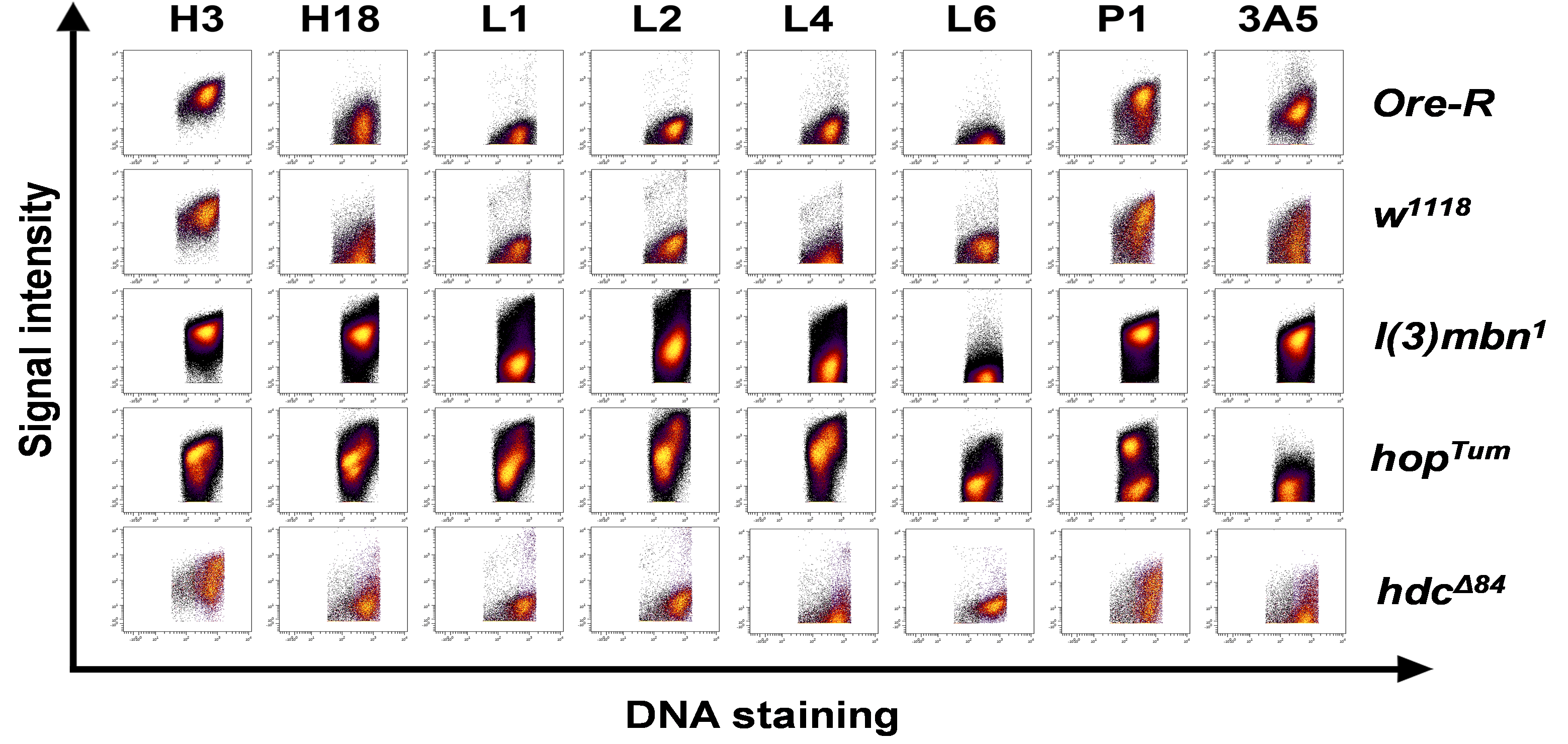

### Figure S7.tif

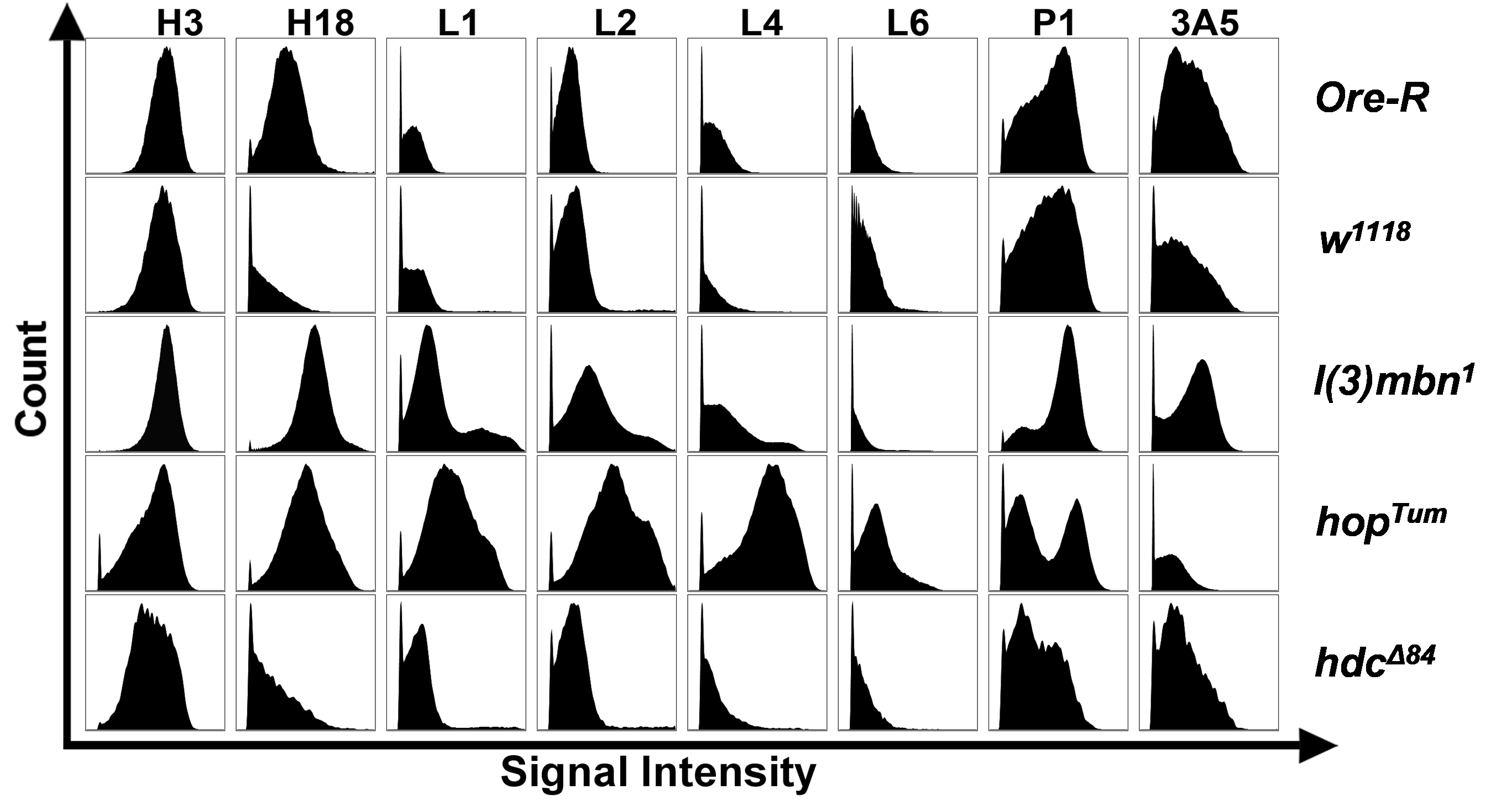

### Figure S8.tif

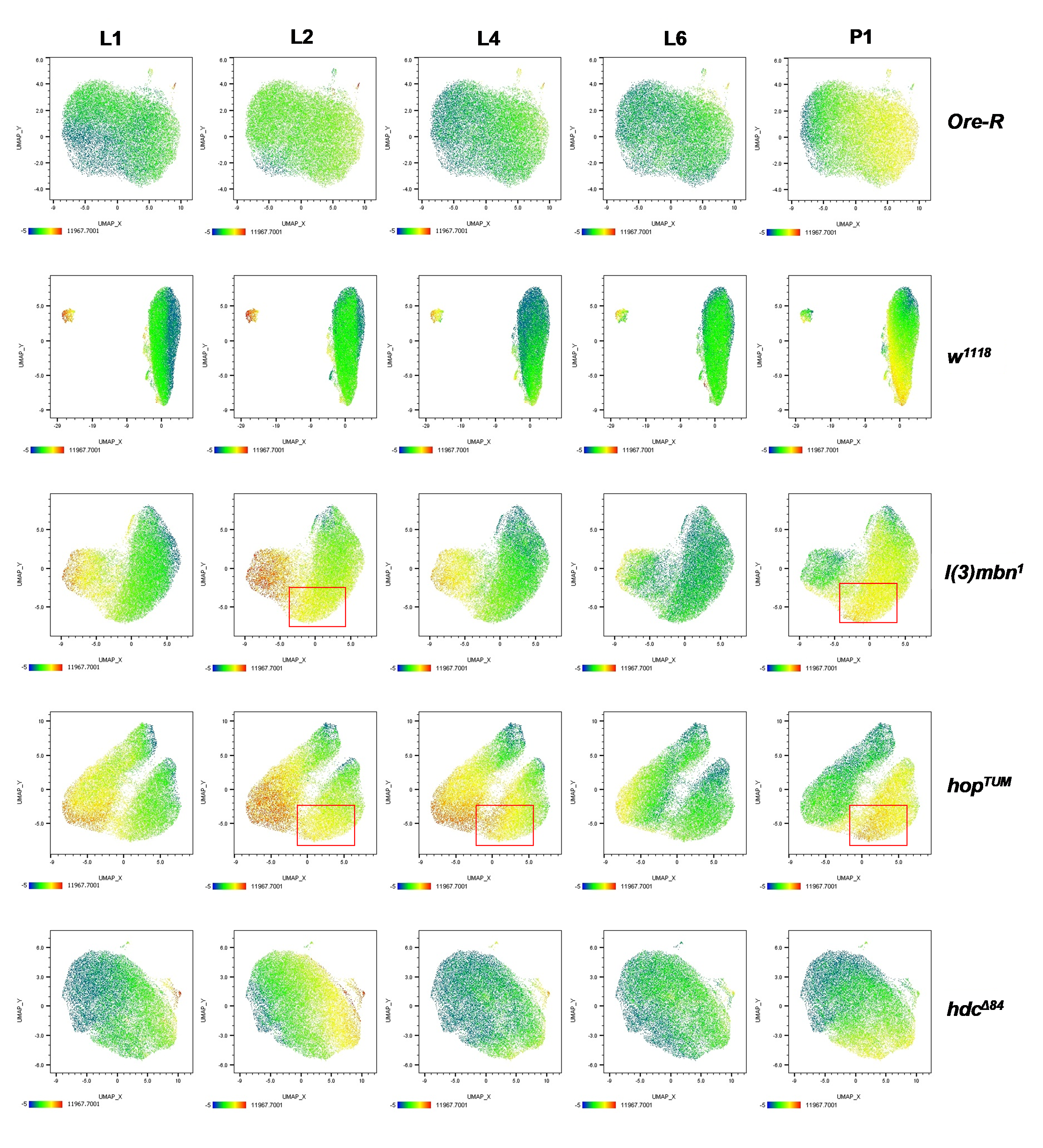

### Figure S9.tif

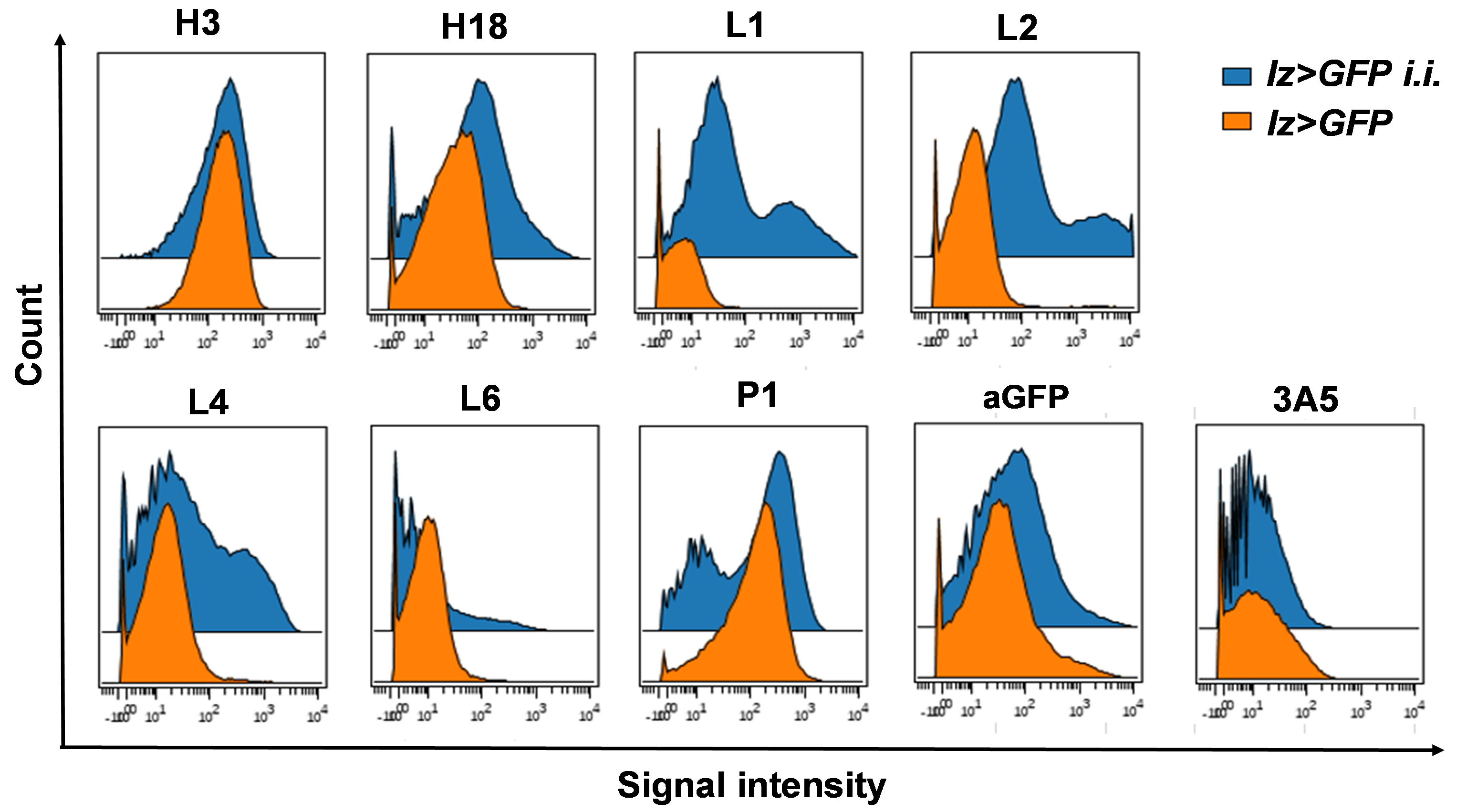
